# Statistical diversity distinguishes global states of consciousness

**DOI:** 10.1101/2023.12.05.570101

**Authors:** Joseph Starkey, Robin L. Carhart-Harris, Andrea Pigorini, Lino Nobili, Adam B. Barrett

## Abstract

Application of complexity measures to neurophysiological time series has seen increased use in recent years to identify neural correlates of global states of consciousness. Lempel-Ziv complexity is currently the de-facto complexity measure used in these investigations. However, by simply counting the number of patterns, this measure theoretically takes its maximum value for data that are completely random. Recently, a measure of ‘statistical complexity’ - which calculates the diversity of statistical interactions - has been devised which aims to account for and remove randomness seen in data. It was recently found that this measure decreases during anaesthesia in fruit flies. This paper investigates this statistical complexity measure on human neurophysiology data from different stages of sleep, and from individuals under the effects of three psychedelic substances: ketamine, lysergic acid diethylamide (LSD), and psilocybin. Results indicate that statistical complexity: (i) differentiates the different stages of sleep analogously to Lempel-Ziv complexity; (ii) increases relative to placebo for all three psychedelic substances. Thus, statistical complexity is a useful alternative measure for investigating the complexity of neural activity associated with different states of consciousness.

## Introduction

A body of recent research has established that several global states of consciousness can be robustly distinguished via measures of temporal differentiation that quantify the number or entropy of temporal patterns in neurophysiological time series (Sarasso et al, 2021), consistent with the so-called ‘entropic brain hypothesis’ (Carhart-Harris et al, 2014; Carhart-Harris, 2018). Time series from unconscious states such as general anaesthesia and non-rapid eye movement (NREM) sleep tend to exhibit less temporal differentiation than time series from awake states (Zhang et al., 2001; Burioka et al, 2005; Casali et al., 2013; Schartner et al., 2015; Andrillon et al, 2016; Schartner et al., 2017b; González et al, 2019), while those from states induced by psychedelics including ketamine, lysergic acid diethylamide (LSD) and psilocybin are associated with increased temporal differentiation (Carhart-Harris et al, 2014; Schartner et al, 2017a; Mediano et al, 2020; Ruffini et al, 2023). The number of patterns is typically computed with versions of Lempel-Ziv (algorithmic) complexity, which counts the number of distinct sequences on binarised or symbolised data (Lempel and Ziv, 1976); entropy measures compute Shannon entropy of variously defined microstates across various temporal scales (Costa, Goldberger and Peng, 2005; von Wegner et al, 2018; Keshmiri, 2020; Mediano, Rosas et al, 2023). These measures are purported to reflect the richness of conscious phenomenology (Carhart-Harris et al, 2014); greater temporal differentiation is associated with richer and more diverse phenomenal content (Schartner et al, 2017b; Tononi and Edelman, 1998), as shown here for example (Timmerman Slater et al, 2023b). However, these measures theoretically take their maximum value for data that are maximally random, i.e. when each data point is completely statistically independent. Thus, there are potentially scenarios in which they do not properly capture the diversity of the actual dynamics, i.e. the diversity of statistical interactions between microstates, but rather unrelated or only tangentially related randomness.

Recently Muñoz et al. (2020) have applied a measure of statistical complexity that does, in theory, capture dynamical diversity. They studied local field potential data from fruit flies (drosophila) and found that statistical complexity decreased in general anaesthesia, compared with an ordinary waking state. The statistical complexity measure calculates the entropy of a time series, but with states specially defined such that if two sequences of observations lead to very similar probability distributions for future observations, then those two sequences are considered identical microstates. In other words, the dynamical complexity is obtained from the minimal model that statistically reproduces similar data, i.e. an epsilon machine (Crutchfield and James, 1989). Statistical complexity is, in theory, zero for maximally random data, when the past tells you nothing about the future-then all states are considered identical. To score high, there must be high signal diversity, but dynamical evolution from distinct sequences needs to mostly be distinct. Broadly, statistical complexity tends to follow an inverted-U shape function of the overall level of randomness, while Lempel-Ziv complexity is monotonically increasing (see Fig. 1).

**Figure 1:**
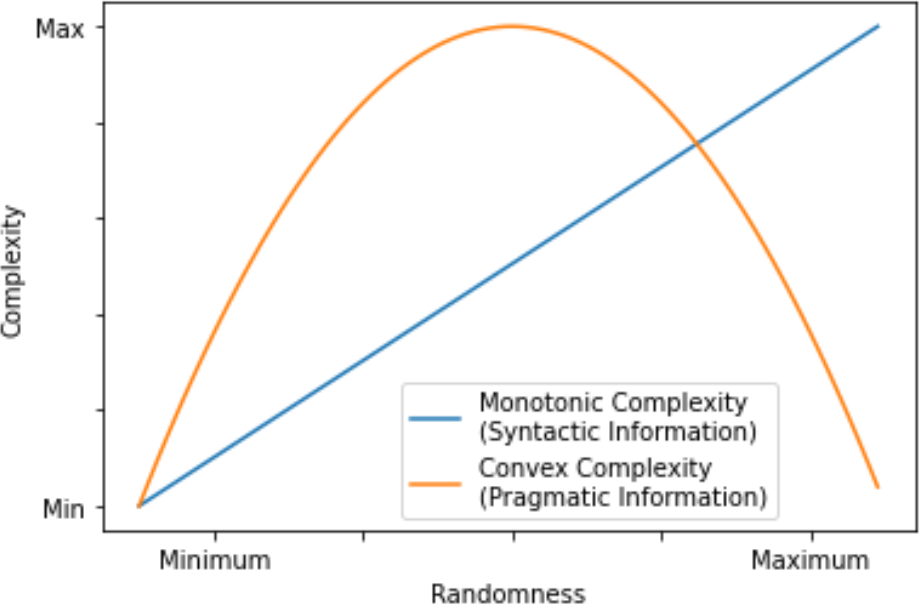
Complexity can have a monotonic or convex relationship with randomness. Lempel-Ziv complexity follows a monotonic “syntactic information” increase, while statistical complexity is designed to follow a convex “pragmatic information” change (Atmanspacher, 2016).

Here we compute statistical complexity on two datasets of human neurophysiological recordings from diverse states of consciousness, on which Lempel-Ziv complexity has previously been analysed (Schartner et al, 2017a; Schartner et al, 2017b): (i) spontaneous depth electrode recordings from epilepsy patients, taken during wakeful rest and different stages of sleep; (ii) magnetoencephalographic (MEG) recordings during altered states of consciousness induced by three psychedelic substances-psilocybin, ketamine and LSD. We analyse and compare the differences in statistical complexity and Lempel-Ziv complexity between the different states of consciousness, when computed on individual channels over time. Our results suggest that NREM sleep generally decreases both measures compared to wakeful rest and REM sleep, while the psychedelic substances increase both measures, most notably LSD.

## Methods and materials

### Data Acquisition

#### Sleep data

The sleep dataset consisted of processed recordings from platinum-iridium semi-flexible multi-contact stereotactically implanted depth multi-lead electrodes of diameter 0.8mm, contact length 1.5mm, inter-contact distance 2mm, ≤ 18 contacts per electrode. These electrodes recorded data from 10 epilepsy patients suffering from ‘drug-resistant, focal epilepsy’, implanted for surgical purposes. Only electrodes outside the epileptogenic zone have been analysed in this study. This means the patients’ epilepsy, theoretically, should not substantially affect the results of this investigation. Four different conditions were identified by an experienced clinician based on polysomnography: i) resting wakefulness (WR), ii) REM sleep, iii) the first stable N-REM sleep episode of the night (NREM1); iv) the final stable NREM sleep episode of the night (NREM2). The two former conditions (WR and REM) were considered “conscious” conditions, as patients were aware of their environment during wakefulness and were likely dreaming during REM sleep, whereas the two latter conditions (NREM1 and NREM2) were considered unconscious states. For each condition, for each participant, there were around 10 minutes of artefact-free recordings from between 18 to 31 channels, recorded at 1000Hz, downsampled to 250Hz, and broken into 10s segments. For full details of the recording and preprocessing, see Schartner et al. (2017b). For standardisation between the two datasets used in this study, a further round of dividing data into 2s segments was carried out, and to each of these segments, linear detrending, baseline subtraction and normalisation of standard deviation was performed. Since the complexity measures we consider take binary data as input, binarisation was carried out on each individual channel for each segment; this was done by thresholding with respect to the median.

#### Psychedelics data

The psychedelics dataset consisted of MEG recordings from a non-invasive CTF 275-channel radial gradiometer system, from a varying number of participants per drug tested. Participants were tested under the effects of three drugs, as well as under a placebo for said drug: 19 participants were tested for ketamine (KET), 15 for LSD, and 14 for psilocybin (PSIL). For each participant, an additional 29 reference channels were recorded for noise cancellation purposes, and each recording was taken at 1200 Hz, later downsampled to 600Hz and divided into 2-second segments, as well as being band-pass filtered with a range of 1-150 Hz. For full details of the recording and preprocessing, see Schartner et al. (2017a). For standardisation between the two datasets used in this study, the 600 Hz 2-second segments were each downsampled to 250 Hz. Similar to the sleep data, linear detrending, baseline subtraction and normalisation of standard deviation was applied to each of these segments, before using the median value to binarise these time series.

### Complexity measures

#### Lempel-Ziv complexity

Lempel-Ziv complexity (LZ) is based on the compressibility of data. It calculates the total number of distinct patterns in a binary string, as demonstrated below in Fig. 2. A string of all 0’s or all 1’s has the minimum possible LZ for a sequence of its length. In contrast, a completely random string will tend to score the highest. Here we normalise LZ scores, by dividing by the maximum possible LZ score for the given length of string (obtained via a random shuffling of the string).

**Figure 2:**
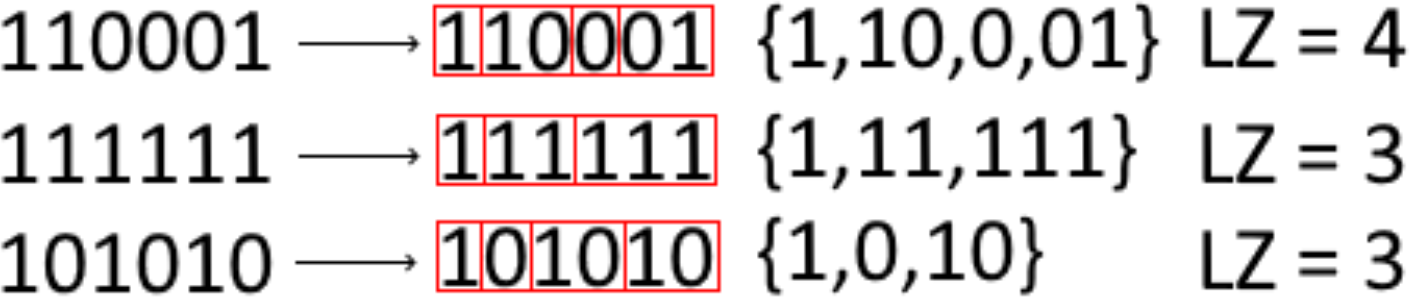
Lempel-Ziv complexity for three example strings. From the start of the string, each character is analysed and added to a “search string”. If this search string is identical to a previously discovered string, the next character in the sequence is added to the search string. If it is not, the search string is added to the list of previously discovered strings, and then the search string is “reset” to an empty string before moving on to the next character in the sequence. This process repeats until the end of the sequence is reached.

#### Statistical Complexity

Statistical complexity is given by the Shannon entropy of an epsilon machine (ε machine), fitted to a time series. An ε machine is essentially a prediction model that most efficiently predicts the future of a time series, assuming that the dependence between past and future states does not change over time (Crutchfield and James, 1989). Possible histories of the system are coarse-grained such that two histories are considered identical if the probability distribution for the future is the same for both the possible histories. The statistical complexity *C*_µ_ is then the standard Shannon entropy over this coarse-graining *S*:

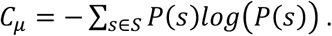

In practice, on finite data, an ε machine cannot be fit exactly, and several hyperparameter choices are required. First, one must choose the length of history to consider, a parameter we call the memory length λ. (If an ε machine is to be accurate at predicting a stochastic process of Markov order *m*, the memory length λ must be greater than or equal to *m*.) Second, for the “probability distribution for the future”, one must choose whether to consider only the next observation, or a string of future observations. Third, a tolerance parameter σ is needed, such that two states are considered the same if their probability distributions of future states are substantially similar, i.e. there is no future state for which the two states disagree in the probability of leading to said future state by more than σ.

As mentioned above, the key feature of statistical complexity is that, in general, it peaks for intermediate levels of randomness, in contrast to LZ, which peaks for maximum randomness (see Fig. 1). Consider the basic example of a binary string, and statistical complexity with the λ parameter set to 1. If 0 is followed by 1 90% of the time, while 1 is followed by 1 70% of the time, then 0 and 1 would be considered distinct states, and the statistical complexity is greater than 0. If, however, 0 is followed by 1 50% of the time and 1 is also followed by 1 50% of the time, then 0 and 1 are considered identical states on the ε machine, and the statistical complexity is 0.

The dataset here consists of time series of length 500 with binary observations. Thus, for memory length λ, the maximum number of states in the ε machine is 2^λ^. So that there is sufficient opportunity for all states to occur a reasonable number of times, the maximum memory length considered was 5. In Supplementary Material (Fig. S2) it is shown that for completely random time series of length 500, the statistical complexity takes close to maximal values when the probability distribution for the future is calculated over strings of length λ (for λ from 1 to 5). Since the whole purpose of considering ε machine based statistical complexity is to have a measure that peaks at intermediate levels of randomness, it was thus decided to calculate the probability distribution for the future based on only the next observation. This led, on the data, to many cases where the number of states in the ε machine was less than the maximum possible, and hence a penalty on excessive randomness. The tolerance parameter σ was set to 0.05, having observed similar patterns of results for ranges of values from 0.01 to 0.1. Statistical complexity was normalised within the results by dividing by a relevant value, utilising the global wakeful rest average at λ = 5 for the sleep study data, or the appropriate placebo result in drug data. Fig. 3 gives an example of calculating statistical complexity for λ = 2.

**Figure 3:**
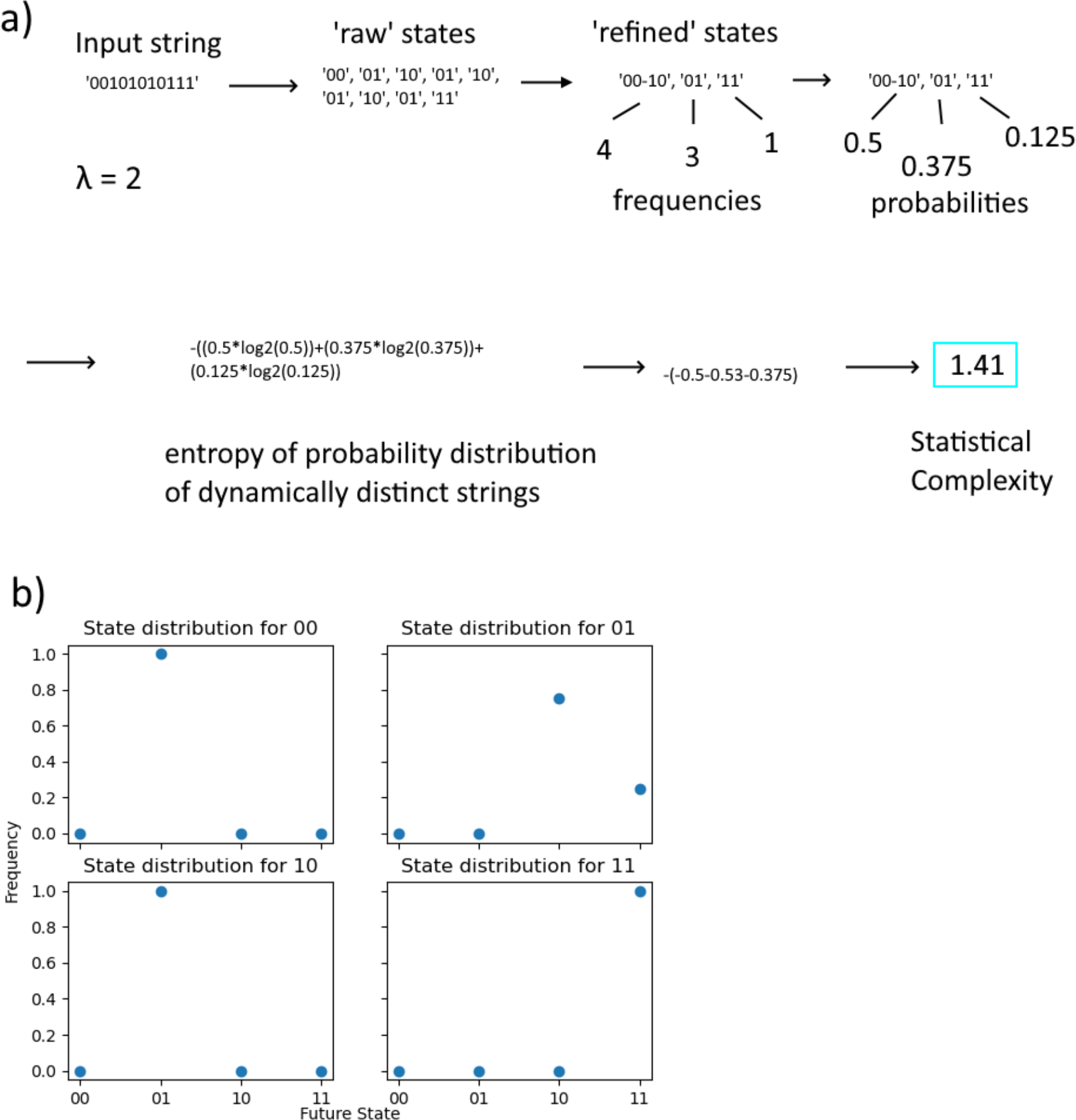
Statistical complexity example calculation (for λ = 2 and σ = 0.05). a) shows that a binary string is broken into strings of length 2, and these are collapsed based on the successors of each string. Strings with sufficiently similar successor distributions are considered the same state. Note that ‘00’ and ‘10’ are considered the same state after refining, a conclusion which can be understood with b). 00 and 10 will always lead to state 01 (i.e. a difference of 0.00, less than σ), and as such are considered the same state for the purposes of calculations.

### Statistics

Group level analyses were conducted via t-tests. At the single participant level, effect size differences between states were measured using Cohen’s d (Cohen, 1988). We call an effect size large if *d*>0.8 and medium if 0.5<*d*<0.8.

## Results

### Sleep data

Mean values over channels were obtained for each complexity measure for each segment, for each state and each participant. These were then averaged across segments to obtain a grand mean for each state for each participant. Fig. 6(a-c) plots the mean and standard error of these. Both Lempel-Ziv complexity and statistical complexity show the trend of wakeful rest (WR) and REM sleep showing the highest complexity, followed by late night deep sleep (lNREM), and finally early night deep sleep (eNREM). eNREM is consistently shown as substantially different to all other stages using both measures (d>0.8, p<0.01), while WR and REM are consistently not substantially different to each other (d<0.5, p>0.05). The difference between measures, in this case, is shown when comparing lNREM to REM and WR: for Lempel-Ziv complexity, both of these differences are not significantly different at p<0.05 but only at p<0.1. For statistical complexity, however, lNREM and WR show a p<0.01 difference, while lNREM and REM show a p<0.05 difference.

At the single participant level, the distribution of complexity across segments was analysed, according to each measure, and effect sizes computed for differences between states, see Fig. 4(f) and 4(g). The same general pattern is observed across both measures, of eNREM being the most dissimilar to other stages, while WR/REM show the most minor differences, but again there are differences regarding specifics. Here, Lempel-Ziv complexity generally showed more substantial differences between states (greater Cohen’s d), although the differences in statistical complexity between eNREM and WR/REM had d>0.8 for all ten participants.

**Figure 4:**
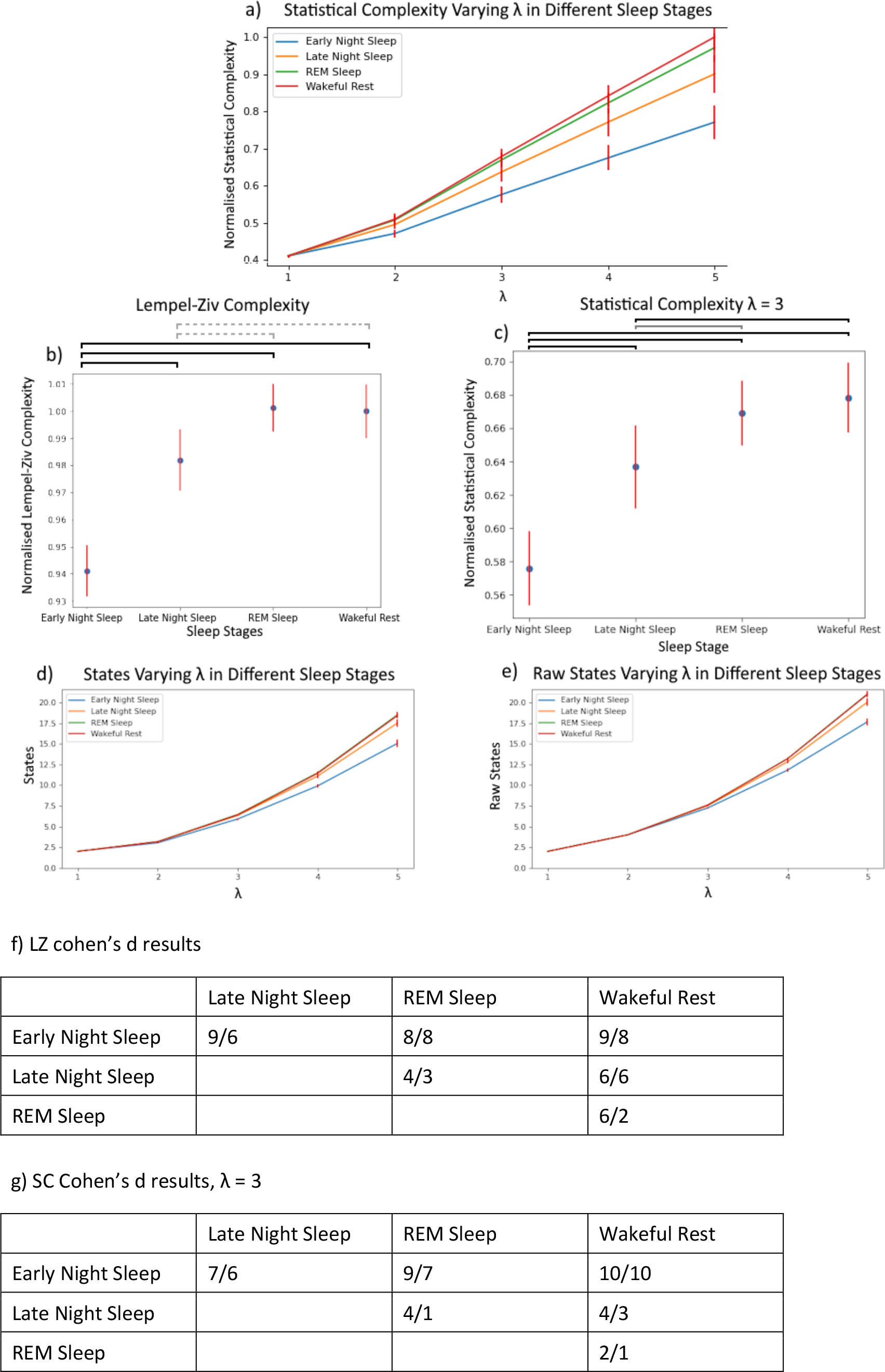
Neurodynamical complexity during sleep. (a) Mean statistical complexity for WR and different sleep stages, for varying λ values. Normalisation is based on mean WR score on a per-participant basis, for λ = 5. (b) shows the results of t-tests for differences between stages using Lempel-Ziv complexity, and (c) shows these for statistical complexity with λ = 3. t-test scores were determined using each participant’s grand mean score as a single data point. These graphs show significant differences at p<0.01 as solid black, p<0.05 as light grey, and p<0.1 as dotted light grey. (d) shows the mean number of states present in the ε machine within a given sleep stage at a given λ, while (e) shows the same pre-’refining’, before states possessing similar future state distributions are merged into single functional states. Full histograms of numbers of states in the ε machine are shown in Fig. S3 in the Supplementary Material. (f) shows the number of participants showing a medium/large (d>0.5/d>0.8) effect size (Cohen, 1988) between two stages using Lempel-Ziv complexity, while (g) shows the same with statistical complexity. In all panels, red bars represent standard error across participants.

The fact that Lempel-Ziv complexity is highest in WR demonstrates that there is greater diversity of patterns in this state. The fact that statistical complexity is also highest in WR suggests this is not entirely due to increased randomness, but also due to increased dynamical diversity (diversity of statistical interactions). To confirm this the number of states in the ε machine was also recorded: Fig. 4(d) shows the mean number of states in the ε machine in each state, and 4(e) the mean number of distinct binary sequences of length λ observed (prior to the merging of dynamically equivalent sequences for the ε machine). These show the same pattern of results as the complexity measures, thus showing that there is indeed greatest dynamical diversity during WR.

In Schartner et al. (2017b), a slightly different version of LZ was used, in which all channels were concatenated before counting the distinct patterns. Thus, there is one value of LZ per segment, in contrast to here, where one value per channel per segment is obtained. Results here are similar to those obtained in Schartner et al. (2017b), but the LZ used here exhibits smaller differences between conditions, when measured with Cohen’s d. In detail, it was found that eNREM and lNREM showed a more substantial difference using Cohen’s d here (6 d>0.8 compared to Schartner’s 2), while lNREM and WR/REM showed a less substantial difference (6/3 compared to Schartner’s 8 for both) as did eNREM and WR/REM (8 for both compared to 10/9).

A regional analysis of statistical complexity was also performed on these data, the results of which can be seen in Supplementary Material (Fig. S1). These broadly mirror the patterns of global results, albeit to differing extents.

### Psychedelics data

Fig. 5(a-c) plots mean statistical complexity for participants in psychedelic states from ketamine (KET), LSD and psilocybin (PSIL), versus placebo. Placebo results are consistently below those of their respective drugs, excepting λ = 1 in which differences are not noticeable. Significance was tested for λ = 3, with a t-test finding p<0.01 for all three drugs (Fig. 5(d)). The results also show that while LSD consistently increases complexity more than KET and PSIL, these latter two drugs still show a significant effect on the group scale. Fig. 5(g) shows the effect size of the difference between drug and placebo per participant. The effect size was large for most participants for LSD, and medium for most participants for KET and PSIL. Regional complexity differences are also shown in Supplementary Material (Table S1).

**Figure 5:**
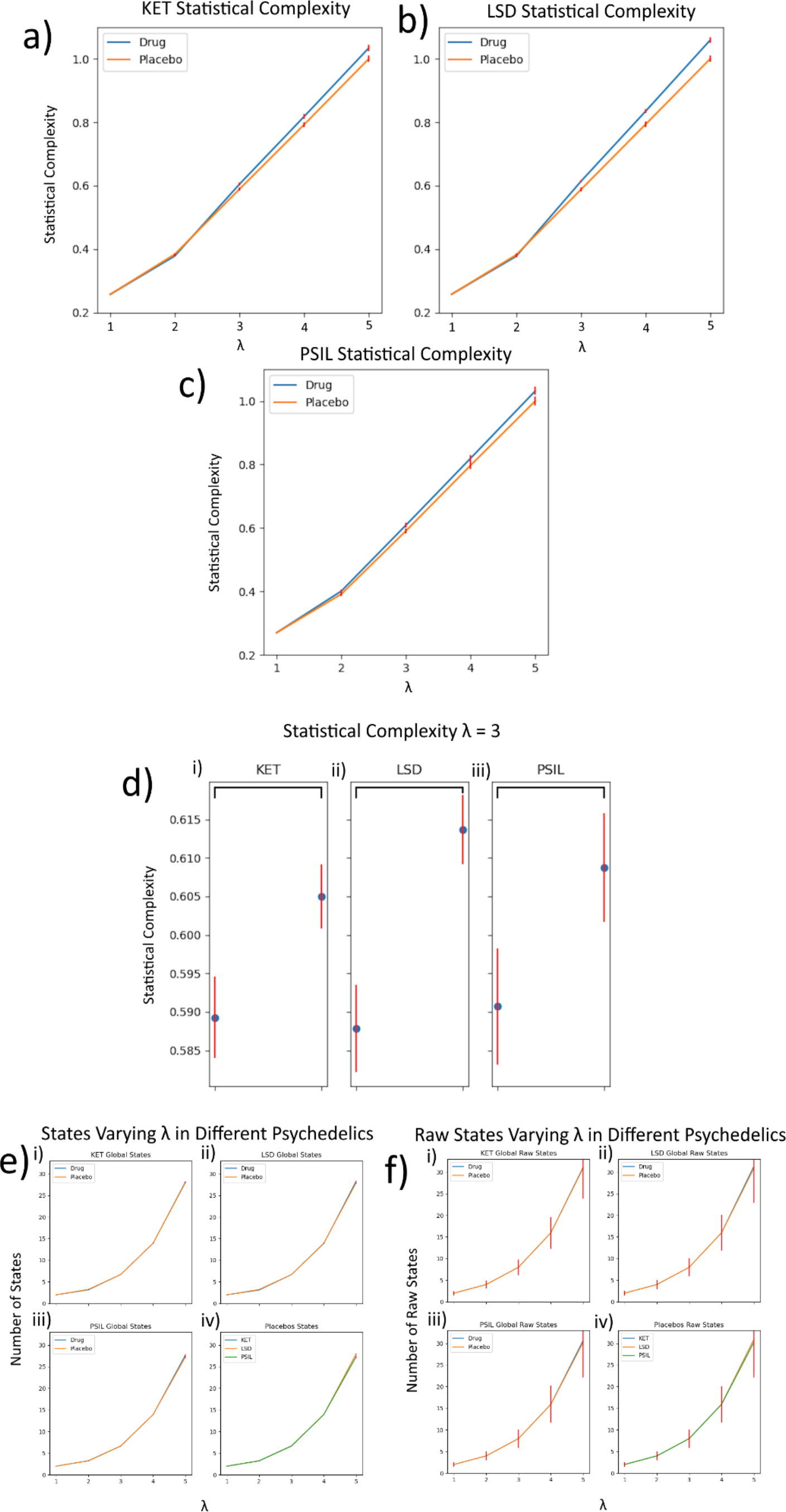
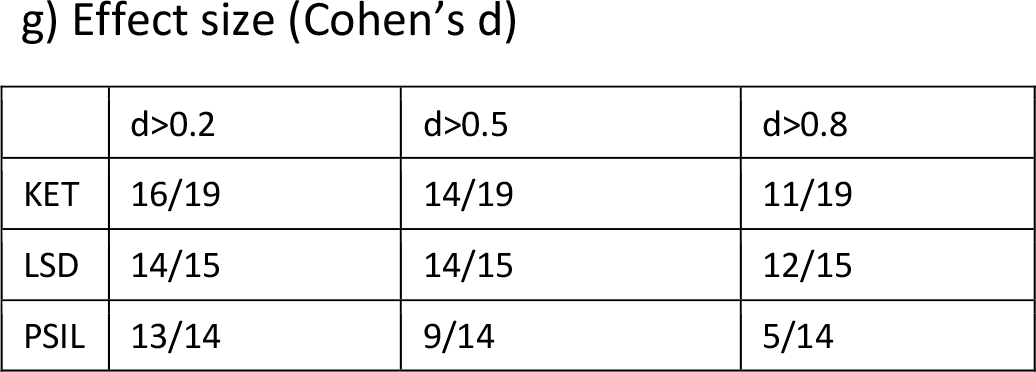
Statistical complexity in the psychedelic state. (a-c) Mean statistical complexity for psychedelics participants with actual drugs versus placebo normalised for KET placebo complexity at λ = 5 equalling 1.0. (d) Mean values for λ = 3, and standard error across participants, black lines signify p<0.01 and grey p<0.05 in t-tests. (e) shows the average number of states present in the ε machine, while (f) shows the same pre-’refining’, before states possessing similar future state distributions are merged into single functional states. (g) The proportion of participants showing a small/medium/large (0.2/0.5/0.8) effect size comparing placebo and non-placebo statistical complexity results according to Cohen’s d. λ = 3 values were used for these calculations. In all figures, red bars represent standard error.

Single channel Lempel-Ziv complexity was already analysed on these data in Schartner et al. (2017a). There it was found that all three psychedelic drugs generally produce an increase in Lempel-Ziv complexity, with LSD and KET generally showing a far more substantial effect that PSIL. The main difference between the results there for Lempel-Ziv complexity, and those here for statistical complexity is that for Lempel-Ziv complexity, PSIL did not produce a significant change in Lempel-Ziv complexity at the population level, when assessed via t-tests, while here statistical complexity increased with significance p<0.01.

The number of states in the ε machine shows no difference between the psychedelic states and the placebos (Fig. 5(e, f)). This suggests that the increase in statistical complexity in the psychedelic state is due to a levelling out of the distribution of times spent in each of the dynamically relevant states (i.e. states of the ε machine).

## Discussion

Results for statistical complexity are broadly consistent with those for Lempel-Ziv complexity on both datasets: namely, relative to wakeful rest, the measure decreases during NREM sleep, is approximately the same during REM sleep, and increases under administration of psychedelic substances. In addition, the number of states in the ε machine reduce during NREM sleep and stay the same in the psychedelic state. Together these results show that the diversity of statistical interactions amongst the different patterns of activity follow the same patterns as the overall diversity of patterns of activity. Thus, it can be concluded that actual dynamical complexity, and not just randomness, decreases in NREM sleep and increases in the psychedelic state.

Compared with Lempel-Ziv complexity, statistical complexity has some extra hyperparameter choices, namely the memory length λ and the tolerance parameter σ. We found similar results for a range of values of σ, but, obviously, if this becomes too small, then it becomes hard to detect states that should be collapsed together in the ε machine, and if it becomes too large, then too many states will be collapsed together. Neural data have long autocorrelations (von Wegner, Tagliazucchi and Laufs, 2017), so it is good to make λ as large as possible, but for limited length time series it must not be too big so that each state has a good chance to occur. Our results found that the effect size grew with λ within the range of values considered. This is a reflection of picking up more details of the dynamics and the increased range of values statistical complexity can take.

Although in theory the statistical complexity is an inverted U-shape function of the level of randomness, in practice, on finite data, it can end up being maximal for maximally random data as seen in the simulations in the Supplementary Material (Fig. S2). Thus, the measure must be supplemented, as done here, with an analysis of the number of states in the ε machine. Since this decreases with increased randomness, this provides vital information about the nature of the dynamical complexity. For both the main results here, this analysis enabled us to conclude that the change in statistical complexity stems from a change in the diversity of statistical interactions and not purely from a change in the overall level of randomness. We have performed simulations of statistical complexity on maximally random time series of the same length as those in the data we analysed. Future work might carry out further simulations to explore more fully the numerical behaviour of statistical complexity with respect to σ, λ, time series length and the level of randomness. Here, following Muñoz (2020) statistical complexity has been computed by sliding along the binary string by a single position rather than λ to isolate states, i.e ‘123456’ would become ‘12, 23, 34, 45, 56’. An alternative to this would be sliding by λ, i.e. generating ‘12, 34, 56’, as this lack of overlap prevents values in the middle of the string being over-represented relative to the ends. However, this substantially reduces the number of observations from which the ε machine can be built, especially at higher values of λ.

The results here add to a body of existing evidence that global states of consciousness can be distinguished by measures of signal diversity (Carhart-Harris, 2018; Sarasso et al, 2021). These results support the entropic brain hypothesis (Carhart-Harris et al, 2014). The entropy/diversity or complexity of spontaneous brain activity that is emphasised in the entropic brain hypothesis is one component of the so-called ‘integrated information theory’ (IIT) of consciousness (Tononi et al, 2016; Albantakis et al, 2023). Specifically, IIT posits statistical information or complexity (referred to by the term ‘information’), along with integration and differentiation between regions (captured by the term ‘integrated’) as key for defining global states of consciousness (Mediano et al, 2022; Tononi and Koch, 2015; Tononi et al, 2016). The entropic brain hypothesis is a well-performing parsimonious model of conscious states – as it shows that conscious states can be described or predicted by just the former component.

A multi-dimensional view of consciousness was proposed by Bayne (2016). However, the adding of dimensions or parameters to a model must be justified as the addition of any new explanatory factor incurs a computational cost. In contrast, elegant models with few parameters aim to explain much with little, which speaks to the value of the entropic brain hypothesis, as endorsed by the present analysis. To our knowledge, a compelling ‘other’ component has not yet been identified that adds appreciable predictive power to the statistical entropy or information component that can justify its computational cost. It is also important to add it does not follow (e.g., it is not an argument of the entropic brain hypothesis) that ‘entropic’ or richer modes of consciousness should entail superior cognitive capacities (Bayne and Carter, 2018). This would be conflating two distinct phenomena: 1) the richness of content of conscious experience – as captured by the entropic brain hypothesis, and 2) cognitive aptitude.

New practical measures of integrated information are still being proposed, but precisely measuring *integration* in the brain remains challenging (Barrett, 2016; Barrett and Mediano, 2019; Hansen and Walker, 2023), and current proposals for measuring integrated information can produce conflicting results (Mediano et al, 2019). As such, well-considered entropy measures remain, to-date, the most powerful approach for predicting conscious states (Carhart-Harris et al, 2014; Carhart-Harris, 2018).

The previous studies by Schartner et al (2017a, 2017b) analysed complexity measures applied to multi-dimensional time series -‘column-concatenated’ LZ complexity, amplitude coalition entropy and synchrony coalition entropy. These also differed with large effect size between sleep stages, but these metrics do not index an integration term – i.e., candidate integration terms as referred to in IIT (Mediano et al, 2022). Simulations have shown that, in practice, all these measures almost entirely capture signal diversity alone, and do not capture any integration phenomenon on finite data (Schartner et al, 2015). Future work will continue to aspire to capture an integration term or phenomenon that can advance on entropic metrics – while justifying the cost of their addition to an explanatory model.

Statistical complexity is more computationally expensive than LZ complexity. For calculating LZ complexity, a string is passed through just once, to obtain the list of patterns that occur. Meanwhile, statistical complexity requires exhaustive analysis of the dynamics following the occurrence of each of the occurring sub-strings. This led to statistical complexity requiring an order of magnitude greater computation time. While not an issue for pure scientific investigation, this perhaps favours LZ complexity for online or ‘real time’ monitoring in potential practical application scenarios, given that LZ complexity distinguishes between the various states equally well as statistical complexity.

Finally, the present study is not the first to propose – and find – that the psychedelic state entails not just more random brain activity, but more *complex*. For example, describing brain function in terms of ‘connectome harmonics’, Atasoy et al (2017) found a greater representation of finer-grained harmonics in the psychedelic state. Further, Luppi et al (2023) found that a broader repertoire of high-frequency harmonic patterns could predict a diversity of conscious states, including the psychedelic state. Similarly, Toker et al (2022) found that an ‘edge-of-chaos’ metric applied to electrocortography (ECoG) and MEG recordings done during diverse states of consciousness could predict conscious ‘level’ in a consistent way as shown here for statistical diversity or complexity. As in the original (Carhart-Harris et al, 2014) and a further iteration of the entropic brain hypothesis (Carhart-Harris 2018), Toker et al (2022) describe their findings as consistent with psychedelics enhancing signatures of system ‘criticality’. This is important, as – like the entropic brain hypothesis – it implies a convex relationship between the entropy of spontaneous brain activity and conscious experience – where, above a critical point, a necessary component for conscious experience may be lost. It could be that an integration term relevant for building a `weak’ IIT (Mediano et al, 2022) is implicitly captured by the Toker et al (2022) metric and the statistical diversity metric validated here. If true, this would imply that IIT could be captured by a single term – implicitly carrying both *information* and e.g., whether this information is structured rather than random. Ongoing theoretical work will assess this.

## Data availability

Code for computing statistical complexity is publicly available at https://github.com/CDR-Clueless/Statistical-Complexity. Data are not publicly available for legal reasons.

## Acknowledgements

We thank Suresh Muthukumaraswamy for giving permission to reuse the psychedelics dataset. JS received funding from the Sussex Centre for Consciousness Science. AP is supported by Progetto Di Ricerca Di Rilevante Interesse Nazionale (PRIN) P2022FMK77. This project/research has received funding from the European Union’s Horizon Europe Programme under the Specific Grant Agreement No. 101147319 (EBRAINS 2.0 Project).

## Notes

### Competing Interest Statement

The authors have declared no competing interest.

