## Supplementary Material for "Statistical diversity distinguishes global states of consciousness"

### 1. Sleep Study Regional Analyses

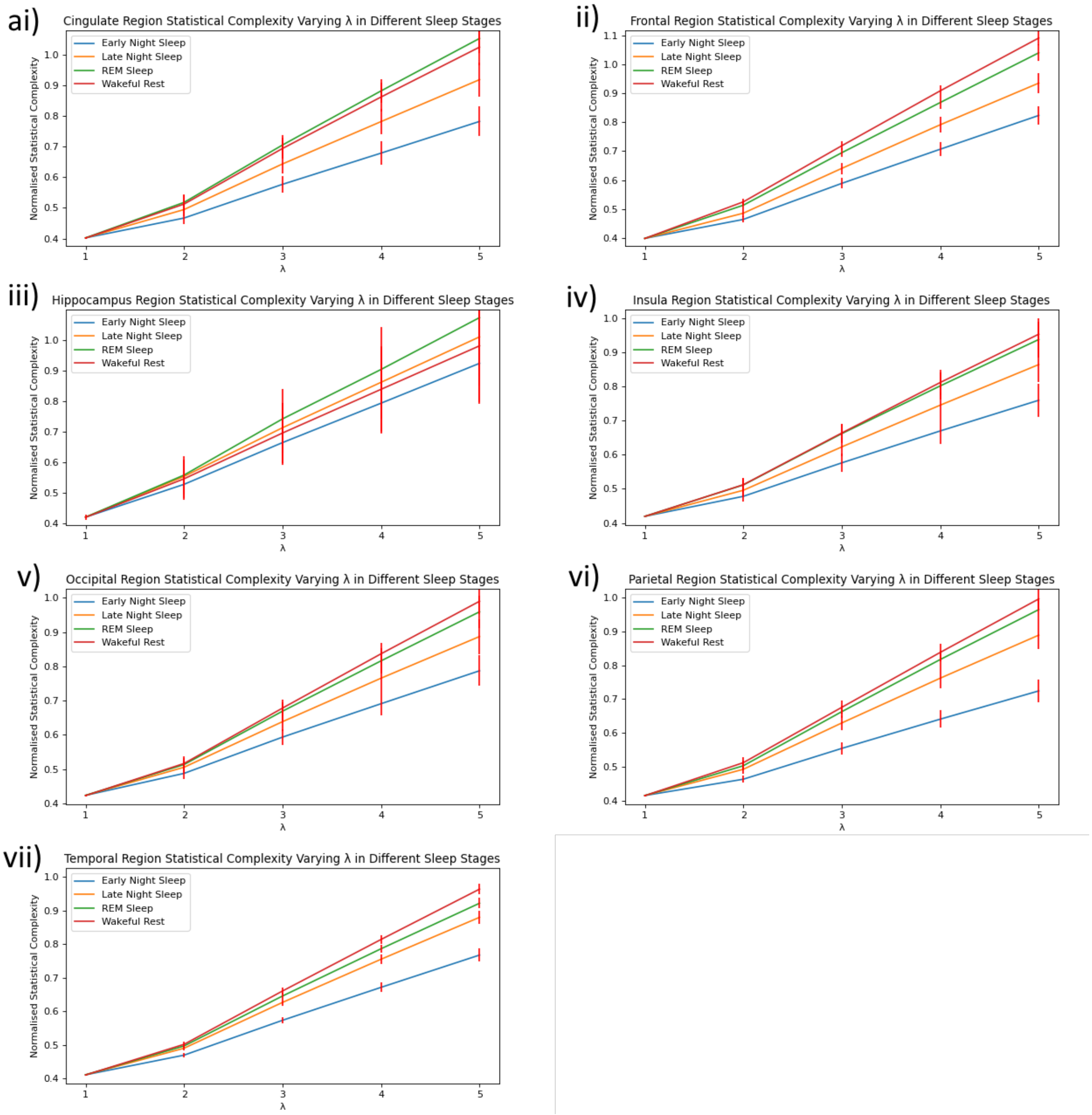

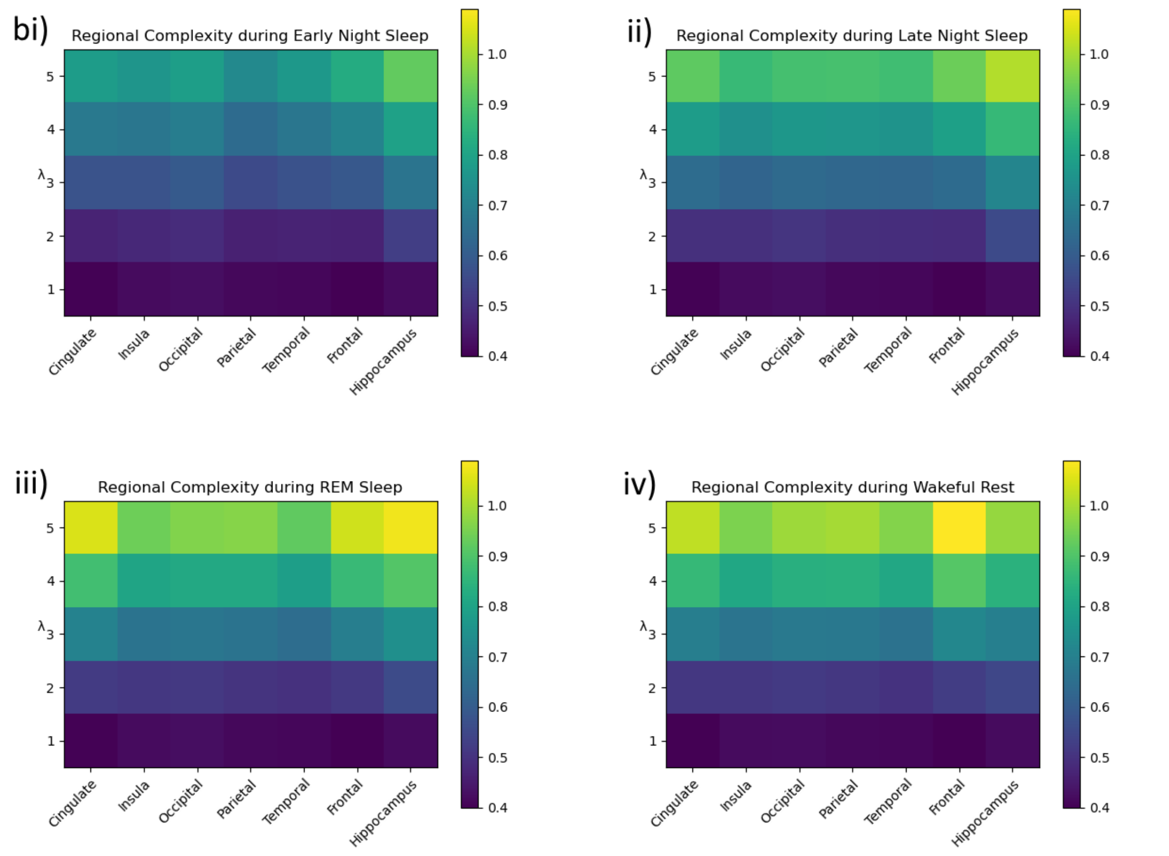


c)

| | Cingulate | Insula | Occipital | Parietal | Temporal | Frontal | Hippocampus |
|---|---|---|---|---|---|---|---|
| WR-REM | 0.19 | 0.20 | 0.22 | 0.21 | 0.23 | 0.22 | 0.20 |
| WR-lNREM | 0.34 | 0.43 | 0.33 | 0.46 | 0.51 | 0.66 | 0.13 |
| WR-eNREM | 0.88 | 0.72 | 0.67 | 1.13 | 1.08 | 1.28 | 0.22 |
| REM-lNREM | 0.44 | 0.43 | 0.23 | 0.38 | 0.48 | 0.62 | 0.13 |
| REM-eNREM | 1.03 | 0.70 | 0.58 | 1.15 | 1.02 | 1.14 | 0.42 |
| lNREM-eNREM | 0.51 | 0.35 | 0.44 | 0.68 | 0.65 | 0.56 | 0.29 |

*Figure S1: Mean statistical complexity in different brain regions and sleep stages displayed as a series of line graphs (a) and heatmaps (b). These results have been normalised so wakeful rest global average at λ=1 equals 1.0, represented through colours labelled to the right. (c) the average Cohen's d scores across participants for each pair of states for each individual region at λ = 3.*

Fig. S1 displays the difference in mean statistical complexity within different sleep stages and brain regions. While the heatmaps in Fig. S1b display the results more concisely, the line graphs of Fig. S1a were shown for easier comparison between sleep stages. These results show the same general pattern as the global results when comparing sleep stages, with a small number of exceptions. Namely, REM sleep shows marginally higher complexity than wakeful rest in the cingulate region, though this difference was found to be insubstantial with Cohen's d as per all WR-

REM differences, and in the hippocampus both REM and lNREM were of higher complexity than wakeful rest, though not to a statistically substantial degree.

#### 2. Psychedelics Regional Analysis

| | Frontal | Sensorimotor | Parietal | Occipital | Temporal | Cingulate | Limbic | Subcortical |
|---|---|---|---|---|---|---|---|---|
| KET | 9/19 | 14/19 | 15/19 | 11/19 | 7/19 | 15/19 | 10/19 | 12/19 |
| LSD | 11/15 | 15/15 | 14/15 | 13/15 | 12/15 | 14/15 | 10/15 | 13/15 |
| PSIL | 3/14 | 7/14 | 9/14 | 9/14 | 9/14 | 10/14 | 11/14 | 9/14 |

*Table S1. Number of participants (out of the total number of participants) showing a large effect size (d>0.8) for difference in statistical complexity between drug and placebo, on a regional basis, for λ=3.*

Table S1 shows the regional statistical complexity results from each psychedelic, in terms of proportion of participants showing a large effect size (d>0.8) difference between drug and placebo. Similar to the global results, LSD consistently shows a more substantial effect on complexity, followed by KET, followed by PSIL (excepting the limbic region, where PSIL affects more participants substantially compared to the other two, and the temporal region, where KET affects less participants substantially.

#### 3. Demonstration that statistical complexity is maximal for fully random data if epsilon machine computed from full future

On finite data, such as the time series of length 500 analysed here, it was considered that statistical complexity may be maximal for random data, due to insufficient sampling of the relevant probability distributions. This was investigated for the case of utilising a memory length λ from 1 to 5, and defining the future as the subsequent λ observations from the present. 100 random binary time series of length 500 were generated and statistical complexity computed on these, see Fig. S2.

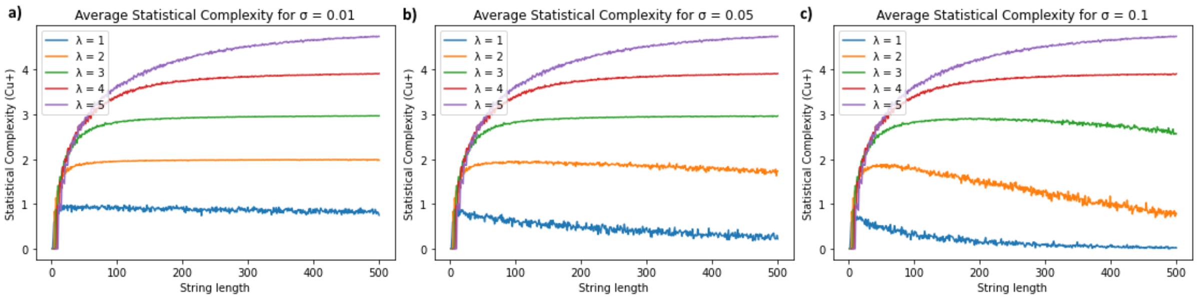


*Figure S2. Average statistical complexity scores of 100 randomly generated samples, with λ = {1, 2, 3, 4, 5} and demonstrating σ = 0.01 (a), σ = 0.05 (b), and σ = 0.1 (c). The maximum possible value of statistical complexity is equal to λ, and this value is approximately what is obtained for time series of length 500 for most of the choices of λ and σ.*

For most choices of memory length λ and tolerance σ, the maximum possible value of statistical complexity is approximately obtained (the maximum value is simply λ, because the maximum number of states in the ε machine is $2^\lambda$ and the maximum entropy of a probability distribution with this number of outcomes is λ). Thus, for the analyses on the real data, the probability distribution for the future was only based on one future observation, to enable the possibility of ε machines with smaller than maximum number of states.

**4. State histograms**

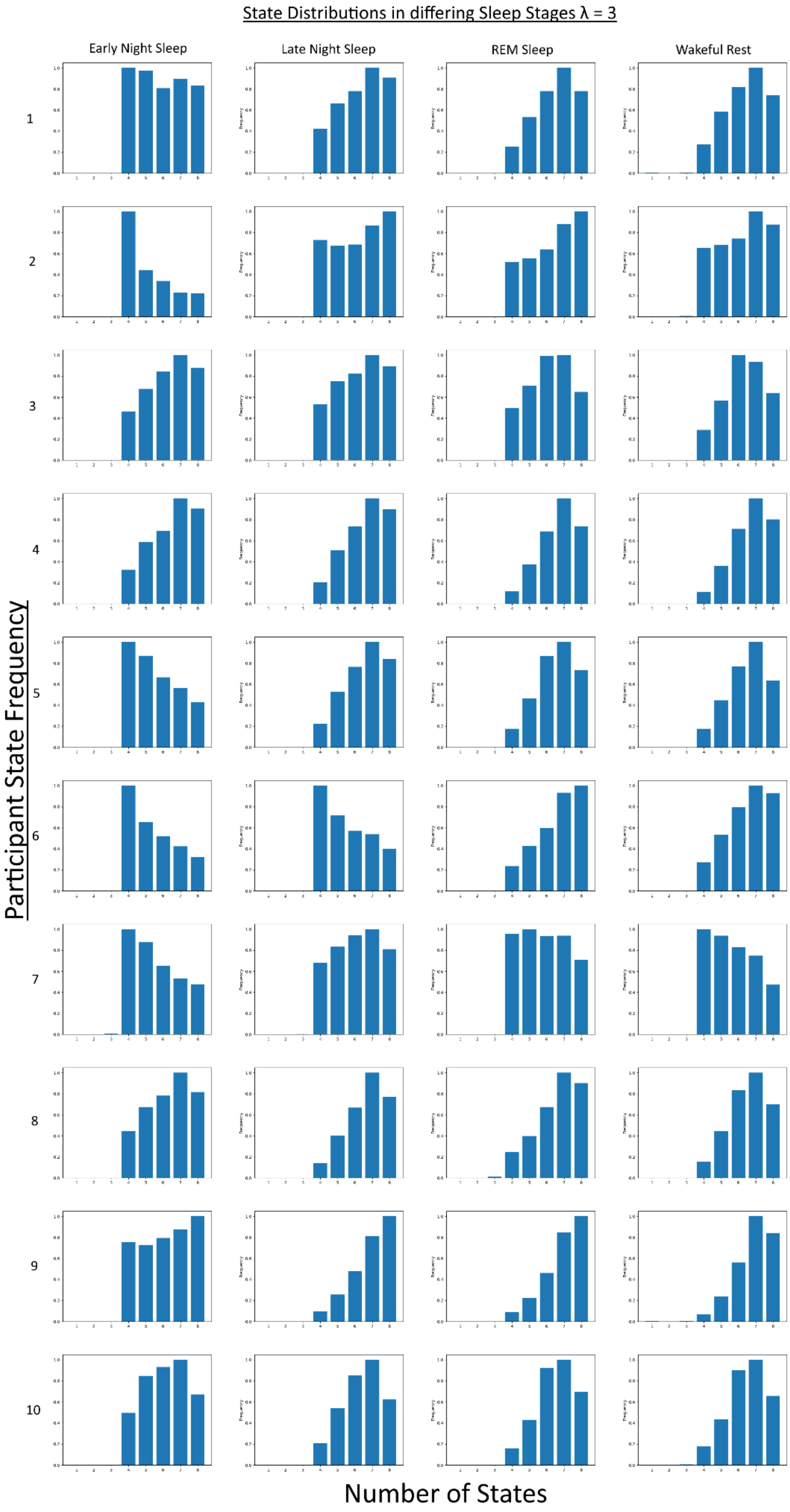


*Figure S3. Distribution across segments of the number of states in the ε machine for λ = 3, per participant for the different states in the sleep dataset.*
